# The M1/M4 agonist xanomeline modulates functional connectivity and NMDAR antagonist-induced changes in the mouse brain

**DOI:** 10.1101/2020.05.01.072595

**Authors:** Caterina Montani, Carola Canella, Adam J. Schwarz, Jennifer Li, Gary Gilmour, Alberto Galbusera, Keith Wafford, Andrew McCarthy, David Shaw, Karen Knitowski, David McKinzie, Alessandro Gozzi, Christian Felder

## Abstract

Cholinergic drugs acting at M1/M4 muscarinic receptors hold promise for the treatment of symptoms associated with brain disorders characterized by cognitive impairment, mood disturbances or psychosis, such as Alzheimer’s disease or schizophrenia. However, the brain-wide functional substrates engaged by muscarinic agonists remain poorly understood. Here we used a combination of pharmacological fMRI (phMRI), resting-state fMRI (rsfMRI) and resting-state quantitative EEG (qEEG) to investigate the effects of a behaviorally-active dose of M1/M4 agonist xanomeline on brain functional activity in the rodent brain. We investigated both the effects of xanomeline *per se* and its modulatory effects on signals elicited by the NMDA-receptor antagonists phencyclidine (PCP) and ketamine. We found that xanomeline induces robust and widespread BOLD signal phMRI amplitude increases and decreased high frequency qEEG spectral activity. rsfMRI mapping in the mouse revealed that xanomeline robustly decreased neocortical and striatal connectivity but induces focal increases in functional connectivity within the nucleus accumbens and basal forebrain. Notably, xanomeline pre-administration robustly attenuated both the cortico-limbic phMRI response and the fronto-hippocampal hyper-connectivity induced by PCP, enhanced PCP-modulated functional connectivity locally within the nucleus accumbens and basal forebrain, and reversed the gamma and high frequency qEEG power increases induced by ketamine. Collectively, these results show that xanomeline robustly induces both cholinergic-like neocortical activation and desynchronization of functional networks in the mammalian brain. These effects could serve as a translatable biomarker for future clinical investigations of muscarinic agents, and bear mechanistic relevance for the putative therapeutic effect of these class of compounds in brain disorders.

## Introduction

Xanomeline is a muscarinic receptor agonist derived from a natural muscarinic agonist, arecoline, an active ingredient of betel nut, chewing of which is common in Asian and Pacific cultures. Of the five muscarinic receptor subtypes, xanomeline exhibits selectivity for the M1 and M4 receptors (Shannon et al., 2000; Andersen et al., 2003; Mirza et al., 2003). An increasing body of preclinical and clinical evidence has highlighted a potential beneficial effect of cholinergic stimulation on brain disorders characterized by cognitive dysfunction or psychosis, including neurodegenerative conditions such as Alzheimer disease or neuropsychiatric diseases such as schizophrenia (Felder et al., 2018; Verma et al., 2018; Erskine et al., 2019). These properties have prompted clinical investigations into the use of this drug to improve cognitive function and behavioral disturbance in Alzheimer’s disease patients, with encouraging results but dose-limiting peripheral cholinergic side effects (Avery et al., 1997; NC et al., 1997; Cui et al., 2008; Si et al., 2010; Wang et al., 2011; Melancon et al., 2013). Interestingly, a parallel set of clinical observations have linked recreational use of betel nut with fewer positive and negative symptoms in schizophrenia (Sullivan et al., 2000), a finding that has been linked to the observation of decreased M1/M4 mAChR density in the brains of schizophrenia patients (for review, see Scarr & Dean 2008).

These anecdotal findings have been corroborated by subsequent research demonstrating putative antipsychotic properties of xanomeline in preclinical and clinical studies. Specifically, treatment with xanomeline has been reported to inhibit the behavioral and motor effects of amphetamine and apomorphine in monkeys (Andersen et al., 2003) and produces behavioral responses in rodents similar to those seen after treatment with traditional antipsychotics (Shannon et al., 1999, 2000; Stanhope et al., 2001). In keeping with these findings, recent clinical studies have demonstrated that monotherapy with xanomeline can improve positive and negative symptoms as well as cognitive function in human subjects with schizophrenia (Shekhar et al., 2008).

While previous studies have addressed neurochemical and behavioral consequences of muscarinic modulation (Mirza et al., 2003; Raedler et al., 2007; Langmead et al., 2008) the macroscopic brain circuits and functional substrates engaged by M1/M4 agonism, and xanomeline in particular, remain poorly investigated. One previous study reported that xanomeline dose-dependently reversed the ketamine-evoked pharmacological fMRI (phMRI) signal increases in the rat brain, demonstrating a modulating effect of muscarinic agonism on an aberrant glutamatergic state considered of mechanistic relevance to schizophrenia (Baker et al., 2012). Since ketamine-challenge phMRI has been successfully translated into healthy humans and modulated by both clinically approved drugs and novel glutamatergic agents (De Simoni et al., 2013a; Doyle et al., 2013; Javitt et al., 2018a, 2018b; Mehta et al., 2018), this observation indicates a potential translational biomarker opportunity. Complementary to this approach, a deeper understanding of the central effects of xanomeline could be obtained by profiling its effect on inter-regional fMRI synchronization, or “functional connectivity”, in distributed brain circuits via resting-state fMRI (rsfMRI) (Errico et al., 2015; Gozzi and Schwarz, 2016). In addition, oxygen (O_2_) amperometry provides a complementary probe-based method to measure the local hemodynamic response in conscious mobile animals (Lowry et al., 1997; Li et al., 2016), and quantitative electro-encephalography (qEEG) provides a means of monitoring the power-frequency profile of the brain’s electrical rhythms more directly, unfiltered by the hemodynamic response underlying the fMRI signals (Ye et al., 2018; Zacharias et al., 2020). Given the established role of cholinergic systems in arousal and neocortical excitation, cholinergic modulation could conceivably also affect large-scale cortical coupling and electrical activity of the brain, resulting in rsfMRI- and qEEG-detectable effects. A characterization of such effects could provide a robust, multi-modal description of the pharmacological properties of this class of compounds of high mechanistic and translational relevance.

Here, we used fMRI, O_2_ amperometry and qEEG to characterize the brain-wide functional substrates modulated by the M1/M4 agonist xanomeline in mice and rats. Specifically, we used pharmacological MRI (Ferrari et al., 2012) to assess the sign and amplitude of the xanomeline-evoked fMRI response, applied resting-state fMRI (rsfMRI) (Sforazzini et al., 2014) to map brain-wide functional connectivity changes produced by central muscarinic stimulation, examined O_2_ amperometric signal changes to corroborate the fMRI effects in conscious moving animals, and used qEEG to profile effects of xanomeline on the frequency profile of the brain’s electrical activity. To mechanistically relate these alterations to the putative anti-psychotic properties of this compound, we also assessed xanomeline’s ability to modulate or prevent brain signals elicited by the non-competitive NMDA receptor antagonists phencyclidine (PCP) and ketamine, compounds that produce robust schizophrenia-like syndromes and stereotypical fMRI, O_2_ amperometric and qEEG responses in rodents (Greene, 2001; Farber, 2003; Gozzi et al., 2008b, 2008c, 2008a; Gass et al., 2013a; Li et al., 2014). Together, these multi-modal experiments provide a comprehensive picture of the effects of xanomeline on brain-wide functional activity, desynchronization of neocortical networks and its modulatory effects on specific NMDAR antagonist-induced network aberrancies. These findings shed light on the large-scale substrates modulated by muscarinic activation and highlight possible translatable biomarkers for future clinical investigations of muscarinic agents.

## Results

### Dose selection

To ensure behavioral relevance of the effects mapped with fMRI, we selected the xanomeline dose based on two independent behavioral assays (Supplementary Figure 1). In the first of these assays (Figure S1A), we measured the ability of xanomeline administration to disrupt conditioned avoidance response. In the second assay (Figure S1B), we measured the effect of xanomeline administration on both spontaneous and PCP-induced locomotor activity. In both assays, for the dose range tested, a dose of 30 mg/kg xanomeline tartrate (i.p. or s.c., respectively) produced maximal efficacy, as demonstrated by complete inhibition of avoidance responses (p < 0.0125, Bonferroni-corrected) and reversal of PCP-induced locomotor activity (F_4,35_ = 17.17, p < 0.0001; Dunnett test, p < 0.05 vs. vehicle). While escape failures during conditioned avoidance testing were not significantly induced by administration of xanomeline tartrate (F_4,36_ = 1.00, p > 0.42), spontaneous activity during the habituation phase of locomotor testing was reduced (F_4, 35_ = 21.34, p < 0.0001, Dunnett test, p < 0.05 vs. vehicle). Based on these results, a dose of 30 mg/kg was selected for testing in imaging studies.

**Figure 1:**
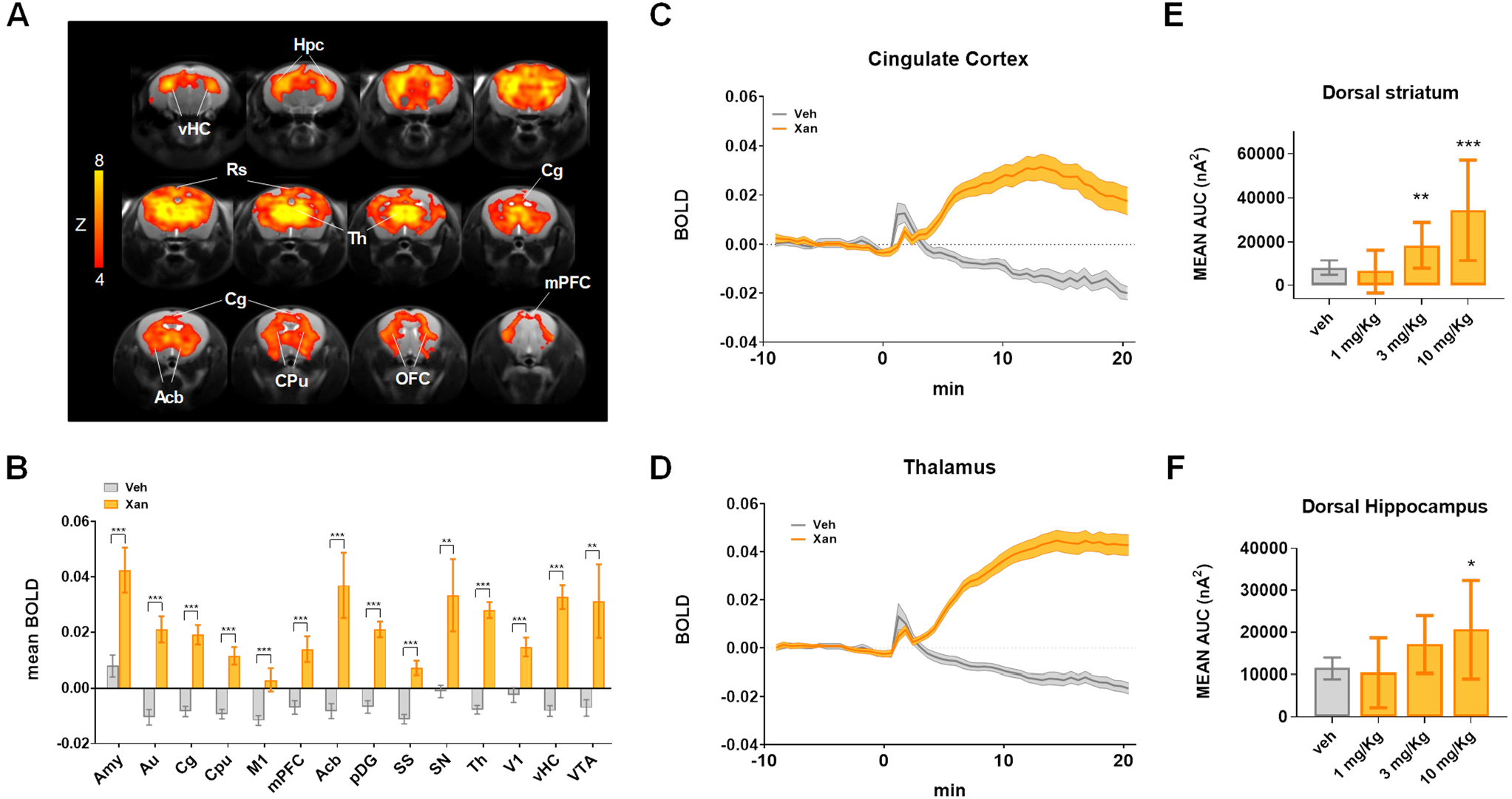
Xanomeline increases functional brain activity as measured with BOLD fMRI and O_2_ amperometry. (A) Maps of BOLD fMRI response elicited by a xanomeline challenge (30 mg/kg s.c.) with respect to vehicle-treated control group. Orange/yellow indicate increases in BOLD signal (Z > 4, p<0.05, cluster corrected). (B) BOLD response to xanomeline in representative ROIs. The effect was plotted as mean BOLD ± SEM. (t-test, *p<0.05, **p<0.01, ***p<0.001, FDR corrected). (C,D) Time course of BOLD signal in cingulate cortex and thalamus, respectively. Xanomeline was injected at time 0. Vehicle n=19, xanomeline n=18. (E,F) O_2_ amperometry changes showing dose-dependent increases measured by probes positioned in the dorsal striatum (E) and dorsal hippocampus (F). Effects are plotted as mean amperometric current ± SEM. (Fishers LSD, *p<0.05, **p<0.01, ***p<0.001, versus Vehicle group). Abbreviation list: Amy, amygdala; Au, auditory cortex; Cg, cingulate cortex; Cpu, caudate putamen M1, primary motor cortex; mPFC, medial prefrontal cortex; Acb, Accumbens; pDG, posterior dentate gyrus; SS, somatosensory cortex; SN, substantia nigra; Th, thalamus; V1, primary visual cortex; vHC, ventral hippocampus; VTA, ventral tegmental area; Hpc, hippocampus; Rs, retrosplenial cortex; OFC, orbitofrontal cortex.

### Xanomeline produces robust brain-wide functional activation

Acute administration of xanomeline produced robust BOLD phMRI responses in widespread brain regions, including retrosplenial, cingulate, medial prefrontal and orbitofrontal cortices, the thalamus, dorsal hippocampal areas, basal ganglia and nucleus accumbens (Figure 1A-D, volume of interest location, Figure S3), with largest effects in the amygdala, accumbens and dopaminergic terminals. The temporal BOLD profile following xanomeline administration revealed a gradual increase in BOLD signal, reaching a maximum in cortical regions approximately 15 min after the injection. Importantly, the peripheral blood pressure (BP) response to xanomeline (Figure S4) showed that drug-induced increases remained within autoregulation range under halothane anesthesia for almost all of the time window imaged (Gozzi et al., 2007), arguing against a non-specific peripheral hemodynamic contribution to the changes observed. To corroborate that analogous findings were also likely to be observed in non-anesthetized animals, we measured the oxygen amperometric response produced by xanomeline in the dorsal striatum and dorsal hippocampus of freely behaving mice (Figure 1E and 1F). Dose-dependent increases in tissue O_2_ levels were observed in both regions (Region x Dose interaction: F_3, 15_ = 4.86, p = 0.0147). In dorsal striatum, both the 3 mg/kg and 10 mg/kg doses of xanomeline significantly increased tissue O_2_ (Fishers LSD: p = 0.0143 and p > 0.0001, respectively) relative to vehicle, while the 10mg/kg dose was significant for the dorsal hippocampus (Fishers LSD: p = 0.0253). Plasma drug level measurements at the end of the fMRI experiments are reported in Table S1. These measurements confirmed successful compound exposure in all animals analyzed in this study.

### Xanomeline decreases local and long-range brain functional connectivity

To map local and long range functional connectivity changes produced by xanomeline we used recently-developed summary metrics described in detail elsewhere (Liska et al., 2016). Xanomeline administration revealed a robust and widespread reduction in both local and long-range connectivity (t>3, p<0.05, cluster corrected, Figure 2A). The effect was especially prominent in cortical regions such as the anterior cingulate, primary motor, auditory, somatosensory and retrosplenial cortices. Interestingly, foci of increased local (but not long range) connectivity were observed in basal forebrain areas and in the nucleus accumbens (Figure 2A). Quantification in areas of interest (Figure 2B,C) corroborated these findings.

**Figure 2:**
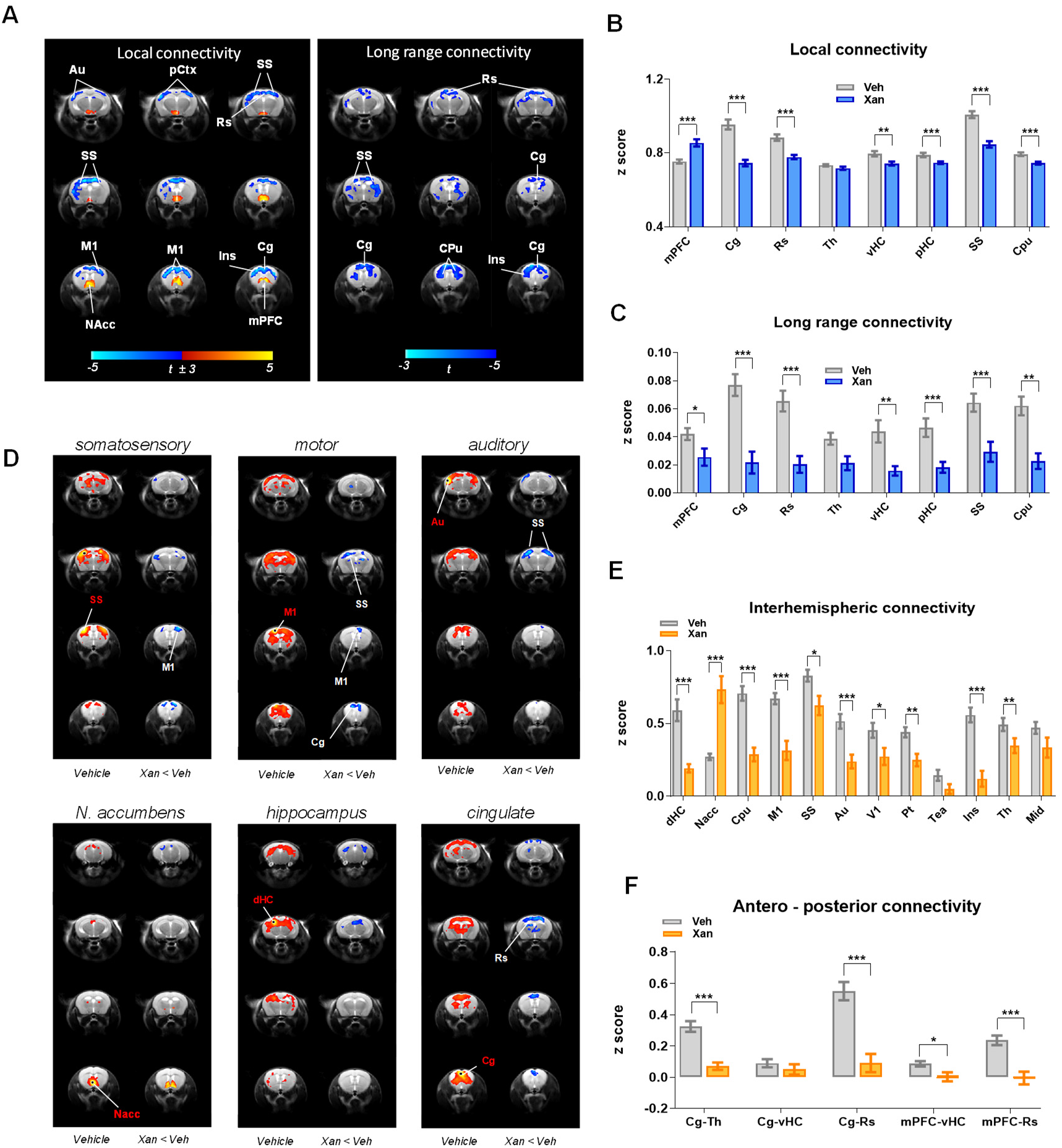
Effect of xanomeline on functional connectivity. (A) Contrast maps showing the difference in local (left) and long-range (right) connectivity between vehicle- and xanomeline-treated animals (one sample t-test, (t>±3, cluster corrected, Veh N=19, Xan, N=18). Blue reflects decreased connectivity, red reflects increased connectivity. (B, C) Quantification of local and long-range connectivity in selected brain volumes (mean ± SEM, student t test, *p<0.05, **p<0.01, ***p<0.001, FDR corrected). (D-F) Xanomeline reduces long-range functional connectivity. (D) Seed-based mapping of selected rsfMRI networks. Network distribution in vehicle-treated controls is shown for reference (Vehicle, N= 19). The effect of Xanomeline (N= 18) is expressed as voxel-wise difference map with respect to vehicle group. Red/yellow indicates correlation with the seed regions (t> ±3, p=0.05, cluster corrected). Blue indicates foci of reduced connectivity in the xanomeline group with respect to control group (t> ±3, p=0.05, cluster corrected). Seed placements are indicated by dots and red lettering. (E) Pairwise functional connectivity between inter-hemispheric volumes of interest. (F) Pairwise functional connectivity between antero-posterior regions of interest. Mean ± SEM, *p<0.05, **p<0.01, ***p<0.001, Student t-test, FDR corrected. Abbreviation list: Au, auditory cortex; pCtx, parietal cortex ; M1, primary motor cortex; NAcc, nucleus Accumbens; Ins, insula; mPFC, medial prefrontal cortex; Cg, cingulate cortex; Rs, retrosplenial cortex; Th, thalamus; vHC, ventral hippocampus; pHC, posterior hippocampus; SS, primary, somatosensory cortex; CPu, striatum; dHC, dorsal hippocampus; V1, primary visual cortex; Pt, posterior parietal association cortex; Tea, temporal association cortex; Mid, midbrain.

### Xanomeline reduces connectivity of inter-hemispheric and antero-posterior networks

To better identify the specific networks underlying these changes in local and long-range connectional deficits, we used seed-based mapping to probe xanomeline modulatory effects in known mouse brain rsfMRI connectivity networks (Figure 2D-F). These analyses revealed the presence of reduced interhemispheric cortico-cortical connectivity in several motor sensory cortical districts (i.e. visual, motor, auditory) as well as in the dorsal hippocampus. Similarly, functional probing of the anterior cingulate cortex revealed reduced antero-posterior functional connectivity across areas of the mouse default mode network (DMN). Interestingly, the nucleus accumbens appeared to be locally hyper-connected, consistent with the increase in the local connectivity parameter described above. A quantification of these effects in region of interests corroborated these effects (Figure 2E-F).

### Xanomeline increases wakefulness and decreases EEG power

To complement the fMRI and amperometric O_2_ measures in the mouse and learn more about the neuronal effects of the drug, xanomeline was also dosed to Wistar rats implanted with EEG electrodes to measure sleep/wake and EEG spectral activity parameters. Drug was administered by sub-cutaneous injection at circadian time −5 (i.e. 5 hours after lights on). At 1, 3 and 10mg/kg xanomeline elicited dose-dependent, robust increases in wakefulness with 37 ± 13 (p = 0.038), 110 ± 13 (p < 0.001) and 225 ± 13 (p < 0.001) minutes of additional wake over the first 7 hours relative to baseline (Figure 3A). At 10mg/kg the increased wake extended into the subsequent dark period, and no rebound sleep occurred as a consequence of the extended wakefulness. Spectral activity occurring during this induced waking was significantly affected with reduction in the alpha and beta range (p<0.001), as well as reduction in the high frequency range (above 80Hz) (p < 0.01 at 3 mg/kg and p<0.001 at 10mg/kg) (Figure 3B-C). Any periods of sleep that were identified in the 7 hours post dosing were significantly depleted in delta power (data not shown).

**Figure 3:**
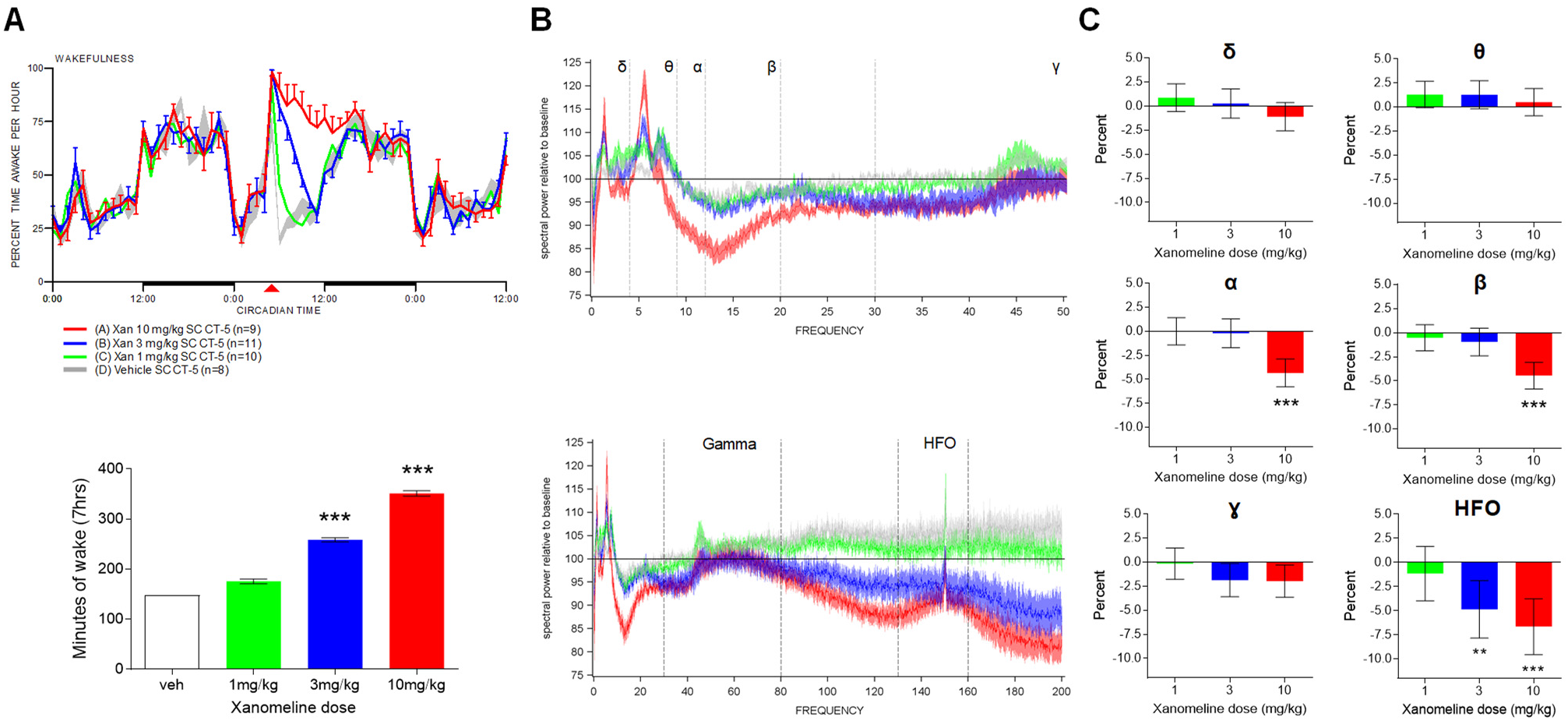
Effect of xanomeline on sleep/wake and EEG spectral activity during wakefulness. Xanomeline increased wakefulness in a dose-dependent manner after s.c. administration. (A, top) Increased wakefulness (percent time awake per hour) was primarily over the first 7 hrs after dosing, returned to normal during the next dark period, and was significant at 3mg/kg and 10mg/kg (A, bottom). Data are displayed as means ± SEM (***p<0.001, one-way ANOVA followed by Dunnett’s test). (B-C) Waking spectral power during this 7 hr period exhibited decreased alpha and beta power (B, top, 0-50 Hz range), unchanged gamma power (B, bottom, full frequency range) and decreased high frequency oscillations (HFO) (B, bottom, full frequency range). Band-specific quantifications in C are expressed as percent change to untreated 24hr baseline (**p<0.01, ***p<0.001, Dunnett’s test).

### Xanomeline attenuates the locomotor and BOLD fMRI response to the NMDAR antagonist PCP

In a subsequent set of studies we assessed the modulatory effect of xanomeline on the functional response elicited by sub-anesthetic dosing of PCP, an NMDAR antagonist with psychotomimetic properties. Acute administration of PCP (1 mg/kg, n=10) induced robust BOLD fMRI activation of a set of cortico–limbo–thalamic regions previously described in the rat and mouse (Figure 4A) (Gozzi et al. 2008; Errico et al., 2015). The temporal profile of PCP-induced activation was similar across activated brain structures (shown for two representative regions in Figure 4B) with a rapid signal increase followed by a sustained activation lasting throughout the imaging window.

**Figure 4:**
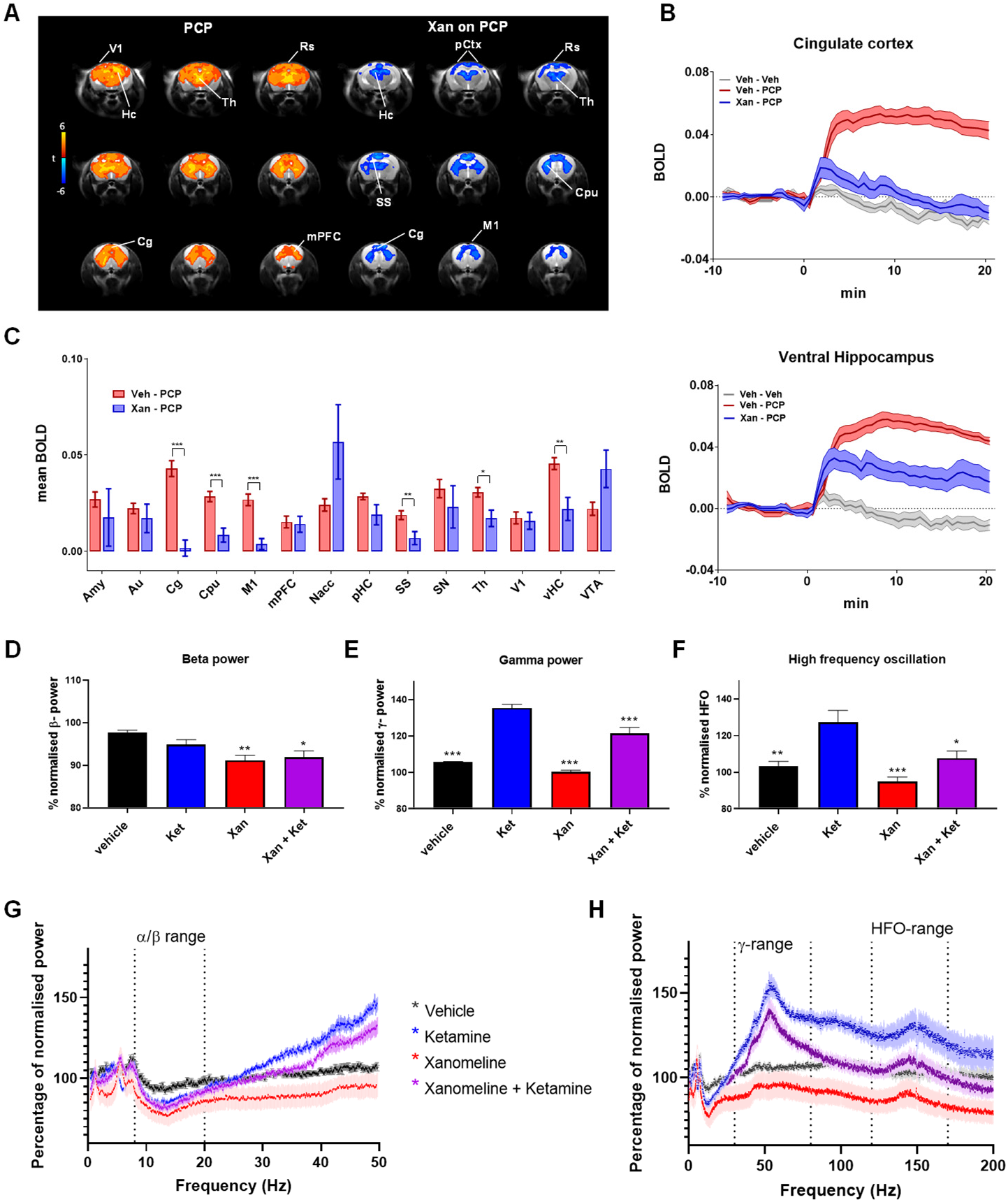
Xanomeline effects on NMDA antagonists. (A) Left: Maps of BOLD response to PCP challenge (1 mg/Kg). Orange/yellow indicate increases in BOLD signal (veh-PCP vs. veh-veh, Z > 4, p<0.05, cluster correction). Right: Contrast maps showing the xanomeline attenuation of PCP-induced fMRI activity (xan-PCP - veh-PCP, Z >3.1, p<0.05, cluster corrected). (B) Time course of BOLD signal in cingulate cortex (top) and ventral hippocampus (bottom). (C) BOLD response as quantified in volumes of interest, veh-PCP n=10, xan-PCP=10, mean ± SEM, *p<0.05, **p<0.01, ***p<0.001, Student t test, FDR corrected (D-H). Xanomeline effects on ketamine-evoked EEG spectral activity. Power change in the EEG spectrum averaged over 6 hrs post dosing with drug or vehicle relative to an equivalent time-period 24 hrs earlier. (D,G) Effects in the 0-50 Hz range with similar drug evoked decreases in the alpha/beta range. (E,F,H) Effects across the full frequency range revealed increased gamma power and HFO with ketamine, and decreases in this range following pretreatment with Xanomeline. Veh (n=13), Ket (n=10), Xan (n=12) Ket + Xan (n=15). Data are displayed as means ± SEM, *p < 0.05; **p < 0.01, ***p < 0.001, one-way ANOVA followed by Dunnett’s test. Significance in (D) is relative to veh, while in (E) and (F) significance is relative to ketamine alone. Abbreviation list: Hc, hippocampus; Cg, cingulate cortex; mPFC, medial prefrontal cortex; Th, thalamus; V1, primary visual cortex; vHC, ventral hippocampus; Rs, retrosplenial cortex; pCtx, parietal cortex; CPu, striatum; Amy, amygdala; Au, auditory cortex; M1, primary motor cortex; SS, somatosensory cortex; SN, substantia nigra; VTA, ventral tegmental area; Nacc, nucleus Accumbens; pHC, posterior hippocampus.

As reported above, behavioral assessments revealed that the dose of xanomeline employed in fMRI studies (30 mg/kg) was able to robustly inhibit the locomotor response produced by acute administration of PCP (Figure S1B). In keeping with these findings, pretreatment with xanomeline robustly attenuated the BOLD response to PCP in across much of the activated areas (anterior cingulate cortex, retrosplenial cortex, primary motor cortex, ventral hippocampus, primary visual cortex, Z>2, cluster corrected) (Figure 4A). When quantified in regions of interest the effect of xanomeline reached statistical significance in most of the examined regions (Figure 4C). Although the response reduction appeared to be rather widespread, complete response blockade was apparent only in selected brain areas, such as cingulate cortex, striatum, and sensorimotor regions.

### Xanomeline decreases ketamine-induced changes in gamma and high frequency EEG activity

NMDA antagonists such as ketamine and PCP have been shown to boost high frequency EEG oscillations with increases in the gamma and high-frequency oscillation (HFO) range (Phillips et al., 2012) and decreases in the beta frequency range (Muthukumaraswamy et al., 2015). To probe whether xanomeline pretreatment could prevent this EEG response, we pretreated Wistar rats with 10mg/kg xanomeline s.c. followed 30 min later by a dose of 10mg/kg (S+)ketamine s.c. at CT-5. Both drugs promoted wakefulness and this was unchanged in the dual-dosed rats (data not shown). The decrease in alpha/beta power was not significantly different with xanomeline, or xanomeline and ketamine, however the ketamine-induced increase in gamma power and particularly the HFO (130-160 Hz) were reduced by pretreatment with xanomeline (Figure 4D-H).

### PCP increases brain functional connectivity in thalamic and fronto-hippocampal networks

We next mapped foci of local and long-range (global) rsfMRI functional connectivity after PCP administration. Voxel-wise mapping revealed focally increased local connectivity in midbrain/collicular regions of PCP-administered animals (Figure 5A, C). A more widespread involvement of sub-cortical areas was apparent in long range (global) connectivity maps, with prominent increases in thalamic and midbrain regions (T>3, cluster corrected, Figure 5A, C-D).

**Figure 5:**
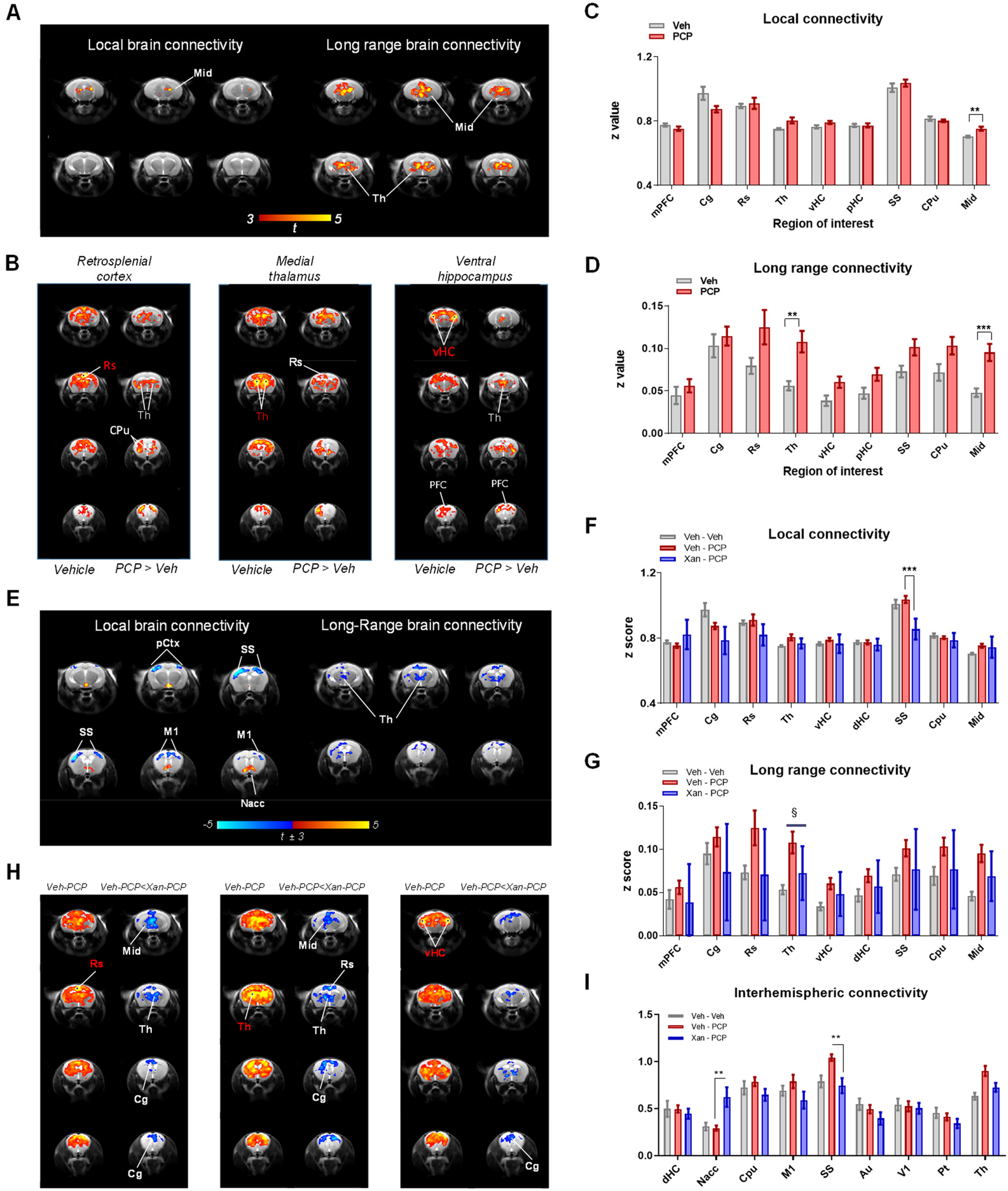
Effect of PCP on functional connectivity, and its antagonization by xanomeline. (A-D) PCP increases brain functional connectivity. (A) Effect of PCP on global and local brain connectivity. Red/yellow reflects increase in connectivity strength (t>3, p<0.05, cluster correction; Veh, N=9; PCP N=10). (B) Seed–correlation mapping highlighted increased connectivity in cortical and subcortical networks of PCP-treated animals. On the left column of each panel, correlation with the seed of interest is reported for vehicle treated animals (n=9; t>2.7, p<0.05, cluster corrected). The right column of each panel indicates the corresponding connectivity difference in PCP-treated animals (yellow indicates connectivity increase in PCP treated group with respect to control, z>2; veh, n=9; PCP, n=10; p<0.05, cluster corrected). (C-D) Quantification of Local connectivity and long-range connectivity in selected brain regions (mean ± SEM, student t test, **p<0.01, FDR corrected). (E-I) Xanomeline attenuates PCP-induced long-range functional connectivity changes. (E) Modulatory effect of xanomeline on PCP-induced local and global connectivity (Xan-PCP vs. veh-PCP, t±3, p<0.05, cluster corrected). (F,G) Quantification of local connectivity and long-range connectivity in volumes of interest (mean ± SEM, student t test, xan-PCP vs. veh-PCP, ***p<0.01, FDR corrected; § p<0.05, uncorrected). (H) Seed-based probing of rsfMRI networks. On the left column of each panel, correlation with the seed of interest is reported for control (vehicle pre-treated) subjects (N=9; t>2.7, p<0.05, cluster corrected). The right column of each panel indicates the corresponding connectivity difference in xanomeline pretreated animals (blue indicates significant connectivity decrease in xanomeline-PCP group (N=10) with respect to control vehicle-PCP, (N=10) Z>2, p<0.05, cluster corrected). (I) Quantification of interhemispheric rsfMRI connectivity (mean ± SEM, student t test, xan-PCP vs. veh-PCP. **p<0.01, FDR corrected). Abbreviation list: Au, auditory cortex; mPFC, medial prefrontal cortex; Cg, cingulate cortex; Rs, retrosplenial cortex; Th, thalamus; vHC, ventral hippocampus; SS, primary, somatosensory cortex; CPu, striatum; Mid, midbrain; pCtx, parietal cortex; M1, primary motor cortex; Nacc, nucleus Accumbens; dHC, dorsal hippocampus. PFC, prefrontal cortex; pHC, posterior hippocampus; V1, primary visual cortex; Pt, posterior parietal association cortex.

To elucidate the brain-wide network substrates affected by PCP, we used seed-based mapping to probe rsfMRI networks previously shown to be modulated by NMDAR antagonism in rodent studies (Gass et al. 2013). Seed-based mapping of polymodal cortical areas (e.g. cingulate, retrosplenial cortices), the thalamus and ventral hippocampal regions highlighted widespread increased functional connectivity in PCP-treated mice (t>2, p<0.05, cluster corrected, Figure 5B). The most prominent effects were associated with the retrosplenial cortex and the thalamus, with PCP increasing connectivity between these substrates and widespread networks of cortical and subcortical areas. Notably, we observed increased connectivity between the ventral hippocampus and the prefrontal cortex, recapitulating similar findings produced by ketamine in the rat (Gass et al., 2013a).

### Xanomeline attenuates PCP-induced long-range connectivity alterations

We next mapped the modulatory effect of xanomeline on PCP-induced local and long-range connectivity alterations by contrasting xanomeline-PCP with vehicle-PCP groups (Figure 5E-G). Consistent with our prior mapping (cf. Fig. 2), xanomeline pretreatment in PCP challenged animals resulted in reduced local connectivity in cortical regions, but focally increased local connectivity in mesolimbic terminals like the nucleus accumbens. Interestingly, contrast maps for global connectivity showed evidence of reduced long range connectivity in thalamic areas in xanomeline pretreated animals, revealing the ability of the drug to prevent PCP-induced increases in thalamic global connectivity. Quantifications of these effect in regions of interest supported this interpretation, showing a reduction in PCP-induced thalamic long-range connectivity (Figure 5G) in animals pretreated with xanomeline (p < 0.05, uncorrected), an effect that however did not survive FDR correction for multiple comparisons. A similar trend was observed in sensorimotor areas and in the retrosplenial cortex.

As above, we next interrogated these functional connectivity effects in specific functional networks using a seed-based approach (Figure 5H-I). This analysis revealed that xanomeline pre-treatment mitigated PCP-induced hyper-connectivity in several brain networks. Specifically, prominent foci of reduced connectivity in xanomeline-pretreated subjects were observed in thalamic and antero-posterior components of the mouse DMN, including fronto-hippocampal circuits. When quantified over large volumes of interest, a normalization of inter-hemispheric somatosensory connectivity was apparent in somatosensory areas, together with an increase in nucleus accumbens coupling in xanomeline treated animals (Figure 5I). Reduced antero-posterior cingulate-retrosplenial and fronto-hippocampal connectivity was also observed in xanomeline pretreated animals (Figure S2), with the former effect surviving FDR correction (q<0.05).

## Discussion

Xanomeline is a functionally-selective M1 and M4 muscarinic agonist that has been investigated as a potential pro-cognitive cholinergic stimulant in Alzheimer disease (Avery et al., 1997; Bodick et al., 1997; Cui et al., 2008; Si et al., 2010; Wang et al., 2011), as well as a putative antipsychotic agent, based on the hypothesis that the muscarinic system is involved in the pathogenesis of schizophrenia (Hyde and Crook, 2001; Lucas-Meunier et al., 2003; Mirza et al., 2003; NR et al., 2003; Sarter, 2005; Raedler et al., 2006; Lisman et al., 2008). The primary original contributions of this report are the demonstration that xanomeline acutely reduces functional connectivity in a widespread set of mouse brain networks and reverses the hyperconnectivity induced by PCP, but focally increasing connectivity in terminals of the mesolimbic pathway in both conditions. Moreover, our findings that xanomeline increases the BOLD signal across several neocortical regions (consistent with a cholinergic stimulant action) and robustly suppresses PCP-induced phMRI-response, generalize previous findings with xanomeline, NMDAR antagonists and their interaction to the mouse. The corroboration of fMRI-derived hemodynamic measurements with EEG recordings of brain activity, amperometric measurements of brain oxygenation as an alternative probe of brain hemodynamics obtained in non-sedated animals, underscores the robustness and substantiates a likely neural origin of the fMRI findings.

Agents that enhance cholinergic transmission have shown potential in improving cognitive function in patients suffering from Alzheimer’s disease (AD) and other memory disorders (Green et al., 2000, 2004). The widespread BOLD signal increase observed with xanomeline in this study is consistent with a robust cholinergic-stimulant effect of the drug in mice, with similarly widespread increased phMRI signal reported in mice undergoing indirect stimulation of basal forebrain cholinergic pathways (Gozzi et al., 2010). Xanomeline dosed 5 hours into the light period of rats produced a pronounced increase in waking state, and a corresponding decrease in NREM and REM sleep as has been observed previously (Gould et al., 2016). Similarly, EEG spectral activity during the waking vigilance state exhibited decreases in the alpha/beta range and while no change in gamma frequency was seen, a dose dependent decrease in high frequency activity (HFO) was observed. However, these wake promoting effects observed in rats do not translate as potentially disruptive to humans at therapeutic doses, as no significant change in normal sleep patterns were observed in Alzheimer’s disease patients (NC et al., 1997).

An earlier study suggested that xanomeline was able to reduce the fMRI increases in BOLD response produced by the NMDA antagonist ketamine in rats (Baker et al., 2012). Here we demonstrate that xanomeline was also able to reduce the increase in high frequency EEG activity produced by ketamine while having no effect on the reduction in lower frequency α/β activity. Interestingly the effects were strongest in the HFO range. NMDA antagonist evoked increases in HFO activity in nucleus accumbens have previously be shown to be reduced by clozapine (Hunt et al., 2015). The ketamine-induced power changes in β −power and γ-power observed here and in other preclinical studies (Phillips et al., 2012) have been recapitulated in humans (Muthukumaraswamy et al., 2015) and correlated with fMRI connectivity changes (Zacharias et al., 2020). However, HFO activity has not yet been reported in humans.

The ability of xanomeline to decrease the functional effects of PCP and ketamine is also consistent with a putative anti-psychotic action of this compound. Numerous human studies have shown that subanesthetic doses of non–competitive antagonists of the NMDA receptor in healthy subjects (Anis et al., 1983), recapitulate cognitive impairments, negative symptoms, and brain functional abnormalities reminiscent of those mapped in schizophrenia (Adler et al., 1999; Coyle et al., 2003; Farber, 2003; Moghaddam and Jackson, 2003; Schwarz et al., 2007). Both the univariate phMRI response and multivariate rsfMRI network activity mapped upon PCP administration are consistent with the result of previous rodent and human mapping of NMDA receptor antagonism, most of which have been carried out with ketamine. Specifically, an increased in cortico-limbo-thalamic fMRI activity was previously shown in rats with PCP, and in humans with ketamine (Schwarz et al., 2007; Deakin et al., 2008; De Simoni et al., 2013b; Doyle et al., 2013; Becker et al., 2016) Moreover, reversal of the ketamine-evoked phMRI response by pre-treatment with two marketed compounds (Doyle et al., 2013) and with novel glutamatergic agents (Mehta et al., 2018) in humans indicates the potential utility of this approach as a translational CNS biomarker. Similarly, a thalamic increase in global (long-range) connectivity, as well as augmented fronto-hippocampal pairwise rsfMRI connectivity have been recently described in rodents and humans (Driesen et al., 2013; Gass et al., 2013b; Anticevic et al., 2015; Joules et al., 2015; Khalili-Mahani et al., 2015). These findings support the translatability of our phMRI/rsfMRI approach across species, and also corroborate the notion that PCP and ketamine produced functionally equivalent fMRI effects, a finding that is consistent with the analogous receptor profile of these two NMDA receptor antagonists.

Xanomeline inhibition of PCP-induced phMRI response is in keeping with the result of a previous fMRI characterization in awake rats (Baker et al., 2012), where xanomeline was shown to dose-dependently suppress the effect of ketamine on the BOLD signal across several regions of interest including association, motor, and sensory cortical regions (Baker et al., 2012). However, establishing these responses in the mouse also opens the way to interrogating brain functional changes in transgenic mouse models such as muscarinic-receptor knock-out mice to establish functional link to specific muscarinic receptor subtypes (Thompson et al., 2018). Moreover, knock-in of humanized muscarinic receptors could avoid dose-response bias due to the different characteristics of rodent and human muscarinic receptors (Chan et al., 2008).

Our data also highlighted a significant widespread decrease in functional connectivity in xanomeline-treated animals. While the electrophysiological correlates of decreased rsfMRI synchronization are unknown, this effect is consistent with an increased cholinergic tone, which increases arousal and leads to neocortical disinhibition at timescales ranging from tens of milliseconds to many seconds (Poorthuis et al., 2014). It is therefore conceivable that the reduced neocortical rsfMRI connectivity may reflect a transition between slow, synchronized oscillations, to desynchronized, stochastic discharge that characterizes the awake/aroused state (Lawrence, 2008). This finding adds to the weight of evidence regarding a possible beneficial action of this drug as a pro-cognitive agent in dementia or neurodegenerative disorders. Interestingly, in the ventro-medial prefrontal cortex and nucleus accumbens, xanomeline produced a local increase in functional connectivity. While the origin and significance of this effect is unclear, it is interesting to note that the drug increases extra-cellular levels of dopamine and functional activation in the same regions (Perry et al., 2001).

The phMRI and rsfMRI data in the present experiments provide strong evidence for a major modulatory role of xanomeline in the central nervous system. The good correspondence between our phMRI results and those obtained with xanomeline and ketamine in awake rats corroborate the notion that light and controlled anesthesia only marginally affects the direction and qualitative patterns of spontaneous or evoked functional activity (see Gozzi & Schwarz 2016 for a recent review). However, a possible interpretational caveat that should be considered when pharmacological manipulations of the cholinergic systems are employed, is a possible direct vascular effect of these drugs. Cholinergic modulators can affect the vasculature directly, with cholinergic agonists causing vasodilatation and cholinergic antagonists vasoconstriction (Sato and Sato, 1995). The administration of acetylcholine in muscarinic M5 knock out mice however did not produce any vasodilatation in the cerebral vasculature, suggesting that the M5 – receptor subtype is responsible for vasoactive functions of acetylcholine in the brain (Yamada et al., 2001, 2003). As xanomeline acts also as a partial agonist at the M5 receptor, a contribution of direct perivascular effects to the readouts mapped in our study cannot be entirely ruled out (Grant and El-Fakahany, 2005), although evidence of the efficacy of xanomeline in counteracting the behavioral (i.e. non hemodynamic) effect of NMDA receptor antagonism MK801 strongly argue against a purely vascular effect of the mapped changes (Barak and Weiner, 2011).

In a recent mouse rsfMRI study, scopolamine, a muscarinic acetylcholine receptor antagonist, caused a functional connectivity disruption in cingulate and prefrontal regions that was recovered by injection of milameline, a muscarinic cholinergic agonist (Shah et al., 2015). It is interesting that these rsfMRI results data are directionally consistent with the functional desynchronization we observed with xanomeline, despite the seemingly divergent pharmacological profile (antagonist vs. agonist) of these two compounds. It is however unclear whether the net resulting effect of scopolamine is functional silencing of activation of cortical cholinergic tone, given the complex pre- and post-synaptic distribution of muscarinic receptors. We also note that oral administration of galantamine, an acetyl cholinesterase inhibitor, produced mostly decreased functional activity in human testing, with areas of functional increase appearing only at a late timepoint (6 hrs post-administration), and mostly in visual regions (Klaassens et al., 2016).

In conclusion, our data document that the M1/M4 receptor-preferring functional agonist xanomeline induces a widespread phMRI response and decreases functional connectivity in the mouse brain. Moreover, xanomeline also reverses the functional and connectivity effects induced by NMDA antagonists ketamine and PCP. These results are consistent with a putative anti-psychotic profile of this drug, and suggest EEG, phMRI and/or rsfMRI as potentially useful pharmacodynamic biomarkers for the determination of central biological activity in human clinical studies.

## Acknowledgments

This study was funded by Eli-Lilly.

## Materials and Methods

### Ethical Statement

All in vivo fMRI studies were conducted in accordance with the Italian law (DL 116, 1992 Ministero della Sanità, Roma) and the recommendations in the Guide for the Care and Use of Laboratory Animals of the National Institutes of Health. Animal research protocols were also reviewed and consented to by the animal care committee of the Istituto Italiano di Tecnologia. All surgical procedures were performed under anesthesia. O_2_ amperometry and EEG experiments were conducted in accordance with the Animals (Scientific Procedures) Act 1986 (UK) following approval from the local Animal Welfare and Ethics Review Body.

### Conditioned Avoidance Responding (CAR)

Male C57Bl/6NTac mice were used (Taconic Farms, Inc., Germantown, NY). The animals were single-housed and maintained in a light- and humidity-controlled room on a 12-h light/dark cycle (lights on at 06:00 h) and behavioral testing occurred during the light phase. Food (Harlan Teklad) and water were available ad libitum in the home cage. Conditioned avoidance responding was conducted in standard mouse shuttle boxes as previously described (Watt et al., 2013). Briefly, mice were first trained to avoid shock at criterion-level performance (≥ 90 % avoidance responding) using the following procedure for two-way active avoidance. Once-daily sessions consisted of a maximum of 50 trials, where each trial began with a CS Phase in which the conditioned stimulus (CS, houselight) was presented and the shuttle door was open. A shuttle response during the CS Phase was recorded as an “avoidance” response. Failure to shuttle during the CS Phase resulted in the US Phase, during which an unconditioned stimulus (US, mild footshock) was initiated. A shuttle response during the US Phase was recorded as an “escape” response. Failure to shuttle during the US Phase was recorded as an “escape failure” response. The occurrence of any one of these 3 response-types terminated the trial. During the intertrial interval (ITI; 30 seconds) the CS and US were not present and the shuttle door was closed. If 20 escape failures occurred in one session, the session was terminated. The percent of avoidance and escape failure responses was calculated relative to the total trials completed per session.

Following reliable demonstration of criterion-level performance, animals were advanced to drug-testing in a within-subjects design where drug-testing occurred approximately twice per week and dose-order per animal was determined using a Latin square (cross-over design). Training (with vehicle) occurred on remaining weekdays, wherein mice were required to qualify via criterion-level performance the day before each drug-testing day. Mice that were unable to achieve or return to criterion-level responding between doses were removed from the study, as were their data. Mice (n = 10, approximately 0.92 yrs. old and 32.8 g) were administered saline and xanomeline tartrate (1, 3, 10, and 30 mg/kg, salt uncorrected, dissolved in 0.9 % saline in a 10-mL/kg dose-volume) via intraperitoneal (i.p.) injection 30 minutes (min) prior to behavioral testing. A one-factor (dose), repeated-measures ANOVA with contrasts against the Saline control was performed using JMP (version 6.0.2) statistical software.

### Spontaneous and PCP-Induced Locomotor Activity (LMA)

Naïve, male ICR outbred mice [Hsd:ICR (CD1^®^)] weighing 21-25 g upon arrival were maintained in a temperature- and humidity-controlled vivarium on a 12:12 light:dark cycle and housed 4 per cage with food (Harlan Teklad) and water available ad libitum. Locomotor activity was conducted in an open field apparatus (Accuscan Instruments, Inc., Columbus, OH), which recorded horizontal photobeam breaks in 5-min increments during the recording window. On the test day, mice received a subcutaneous (s.c.) dose of saline (vehicle) or xanomeline tartrate (1, 3, 10, or 30 mg/kg; salt uncorrected; water vehicle; 10 mL/kg) and were then placed into the open field for a 30-min habituation phase to measure spontaneous locomotor activity. Immediately following completion of the habituation phase, 5 mg/kg phencyclidine HCl (PCP; salt uncorrected; water vehicle) was administered (s.c.) to all mice, which were then returned to the open field for the 60-min PCP-induced activity phase. Total horizontal activity counts for each phase were analyzed using one-way Analysis of Variance (ANOVA) and Dunnett’s test, using the vehicle-treated group (prior to PCP-treatment) as the control for the habituation phase (spontaneous activity) and the vehicle-treated group (after PCP-treatment) as the control for the PCP-induced activity phase. Statistical analyses were performed in JMP v. 8.0.2. On test day, mice received a subcutaneous (s.c.) dose of a saline vehicle or xanomeline tartrate (1, 3, 10, or 30 mg/kg; salt uncorrected; water vehicle) and then placed into an open field apparatus (Accuscan Instruments, Inc, Columbus, OH) for a 30 min period.

### Experimental design for fMRI

Male 12-week old C57Bl6/J mice were purchased from Jackson Laboratories (Bar Harbor, ME, USA). The animals were housed with temperature maintained at 21 ± 1°C and humidity at 60 ± 10%. Two consecutive 30-minute BOLD fMRI time series were acquired; during the first, either vehicle (saline) or xanomeline (30 mg/kg, salt uncorrected) was administered subcutaneously (s.c.) after 10 minutes; during the second, either PCP (1 mg/kg) or vehicle (saline) was administered intravenously (i.v.) after 10 minutes.

Mice were randomly assigned to one of the following treatment groups:

- Group 1: subcutaneous administration of xanomeline (30 mg/Kg) followed by an intravenous challenge with PCP (1 mg/Kg) 30 min later (n=10);
- Group 2: subcutaneous administration of xanomeline (30 mg/Kg) followed by an intravenous challenge with vehicle 30 min later (n=10);
- Group 3: subcutaneous administration of vehicle followed by an intravenous challenge with PCP (1 mg/Kg) 30 min later (n=10);
- Group 4: subcutaneous administration of vehicle followed by an intravenous challenge with vehicle 30 min lat<er (n=10);

The dose of xanomeline was chosen based on the results of the CAR and LMA behavioral assays described above.

#### Data acquisition

The animal preparation protocol was recently described in detail (Ferrari et al., 2012; Sforazzini et al., 2014; Zhan et al., 2014). Briefly, mice were anaesthetized with isoflurane (5% induction), intubated and artificially ventilated (2% maintenance). The left femoral artery was cannulated for continuous blood pressure monitoring and blood sampling. At the end of surgery, isoflurane was discontinued and substituted with halothane (0.75%). Functional data acquisition commenced 45 min after isoflurane cessation. Mean arterial blood pressure was recorded throughout the imaging sessions. Arterial blood gases (p_a_CO_2_ and p_a_O_2_) were measured at the end of the functional time series to exclude non-physiological conditions. Three animals were discarded from the study for physiological reasons. Specifically, one subject exhibited unstable MABP and ventilation problems, two subjects exhibited non-physiological blood gas levels (pO_2_ < 100 mmHg). A statistical comparison of pCO_2_ values using Student’s two sample t-test did not reveal any significant difference between all pairs of treatment groups (p>0.51, all groups).

MRI data were acquired with a 7.0 Tesla MRI scanner (Bruker Biospin, Milan) as previously described, using a 72 mm birdcage transmit coil, and a four-channel solenoid coil for signal reception. For each session, high-resolution anatomical images were acquired with a fast spin echo sequence (repetition time (TR)/echo time (TE) 1200/15 ms, matrix 192 × 192, field of view 2 × 2 cm2, 18 coronal slices, slice thickness 0.60 mm). Co-centred single-shot BOLD EPI time series were acquired using an echo planar imaging sequence with the following parameters: TR/TE 1200/15 ms, flip angle 30°, matrix 100 × 100, field of view 2 × 2 cm^2^, 18 coronal slices, slice thickness 0.60 mm. Two consecutive scans encompassing 1600 volumes each (total duration 32 minutes) were acquired to keep the size of rsfMRI scans manageable and reduce the risk of system failures. Xanomeline or vehicle was injected 12 min after the beginning of the first scan (volume #600). PCP or vehicle was injected 12 min after the beginning of the second functional scan (volume #600).

#### Image analysis

##### Pharmacological MRI

The phMRI response was mapped and quantified as previously described (Errico et al., 2015). Briefly, phMRI time series were spatially normalized to a common reference space and signal intensity changes were converted into fractional BOLD changes. The first 10 volumes of the phMRI time series were removed to permit signal equilibration. After that, individual BOLD EPI volumes were averaged into 36 second (30 volumes) bins to decrease data dimensionality. This sampling proved to be sufficient to describe the slow rising phMRI response to xanomeline or PCP. Voxel–wise group statistics was performed using FEAT Version 5.63, on spatially smoothed data (full width at half maximum of 0.6 mm), using a boxcar function. Group comparisons (treatment vs. vehicle) were carried out using multilevel Bayesian inference, a Z-threshold > 4 and cluster-corrected significance threshold of p=0.05.

##### rsfMRI Image analysis

The first 50 volumes of the rsfMRI data were removed to permit signal equilibration. The time series were then despiked, corrected for motion and spatially normalized to an in-house mouse brain template as previously described (Sforazzini et al., 2014; Liska et al., 2015). Spatially-normalized data had a spatial resolution of 0.1042 × 0.1042 × 0.5 mm^3^ (192 × 192 × 24 matrix). Head motion traces and mean ventricular signal (averaged rsfMRI time course within an average reference ventricular mask) were regressed out of each of the time series. All rsfMRI time series were spatially smoothed (full width at half maximum of 0.6 mm) and band-pass filtered to a frequency window of 0.01-0.1 Hz.

To obtain an unbiased identification of the brain regions exhibiting differences given by the treatment (xanomeline or PCP) in functional connectivity, we calculated long range connectivity (GLBC, global connectivity minus local connectivity) and local brain connectivity (LBC) maps for all subjects (Cole et al., 2010; Liska et al., 2015). This metric considers connectivity of a given voxel to all other voxels simultaneously by computing average connectivity strength. Specifically, we employed the weighted LBC and GLBC method, in which individual r-scores are first transformed to z-scores using Fisher’s r-to-z transform and then averaged to yield the final GLBC voxel value. Local connectivity strength was mapped by limiting this measurement to connections within a 0.6252 mm (6 voxels in-plane) sphere around each voxel, while long-range connectivity was computed by considering only connections to voxels outside this sphere. Voxelwise inter-group differences in each of these parameters were mapped using a Student’s t-test (p < 0.05 followed by a cluster correction with p_c_ < 0.05 as implemented in FSL). The effect was also quantified in atlas-defined volumes of interest (VOIs).

The anatomical location of the examined VOIs is reported in figure S3. Region identification and naming follow classic neuroanatomical labelling described in (Paxinos and Franklin, 2003). Inter-group differences in the extent and intensity of long-range rsfMRI correlation networks were mapped using seed-based approach as previously described (Sforazzini et al., 2014). Small a priori seed regions of 8×8×1 voxels (approx. 0.35 mm^3^) were chosen to cover antero-posterior cortical networks, and representative heteromodal cortical structures (Figure S3). The mean time courses from the unilateral (medial) or bilateral seeds were used as regressors for each voxel. Group level differences in connectivity distributions were mapped using two-sample Student’s t-tests (p < 0.05, cluster correction with p_c_ < 0.05). Alterations in inter-hemispheric functional connectivity were assessed by computing correlation coefficients of inter-hemispheric VOI pairs depicted in Figure S3. The functional connectivity effects of xanomeline or PCP were analyzed over a 20 minute time-window (1000 volumes) following drug injection, capturing the peak of pharmacological effect produced by these two agents.

### Plasma xanomeline/PCP Quantification

Plasma xanomeline quantification was performed in mice from groups 1 and 2; plasma PCP quantification was performed in mice from groups 1 and 3, as previously described (Bertani et al., 2010). Briefly, 100 – 200 ml blood samples were taken from the cannulated femoral artery at the end of the experiment. Plasma was obtained by centrifugation at 5000 g for 10 min at 4 °C and split into aliquots in the presence of a protease inhibitor mix. Samples were stored at −20 °C until quantification, which was performed by Eli Lilly Inc., Indianapolis.

### In vivo O_2_ amperometry

22 male C57BLJ/6NTac mice weighing 21-28g (3 months old) were housed in standard housing conditions (4 per IVC cage, 07:00 to 19:00 light phase, constant temperature and humidity, ad lib food and water) and surgically prepared at Eli Lilly (Windlesham, UK). Changes in extracellular tissue [O_2_] were measured using constant potential amperometry with carbon paste electrodes (Lowry et al., 1997). A potential of −650 mV was applied to electrodes allowing the electrochemical reduction of dissolved O_2_ at their tip (Lowry et al, 1996; Bolger et al 2011). Changes in the measured current produced by the electrochemical reduction of O2 are directly proportional to the local extracellular tissue [O_2_] (Hitchman, 1978). Carbon paste electrodes (CPEs) were constructed from 8T (200μm bare diameter, 270μm coated diameter) Teflon®-coated silver wire (Advent Research Materials, Suffolk, UK). The Teflon insulation was slid along the wire to create an approximately 2-mm deep cavity, which was packed with carbon paste (prepared according to O’Neill et al., 1982). Prior to implantation, all CPEs were calibrated in vitro in a glass cell containing 15 ml phosphate buffer solution (0.01 M), pH 7.4, saturated with nitrogen (N_2_) gas, atmospheric air (from a RENA air pump), or pure O_2_ (compressed gas) at room temperature. The concentrations of dissolved O_2_ were taken as 0μM (N_2_-saturated), 240μM (air-saturated) (Foster et al., 1993), and 1,260μM (O_2_-saturated) (Bourdillon et al., 1982), respectively. Reference and auxiliary electrodes were also prepared from 8T Teflon^®^-coated silver wire by removing 2mm Teflon from the tip. All electrodes were soldered to gold connectors, which were cemented into six-pin plastic sockets (both from Plastic One, Roanoke, VA) during surgery.

Under isoflurane anesthesia, animals were implanted with carbon paste electrodes (CPEs) in the dorsal striatum (AP +0.5 mm, ML-2.2 mm, DV −2.8 mm from the bregma), and the dorsal hippocampus (AP - 1.7 mm, ML- 1.2 mm, DV −1.6 mm from bregma). The reference electrode was placed in posterior cortex, and the auxiliary electrode was wrapped around one posterior skull screw. Pre- and post-operative Rimadyl (Carprofen 5mg/kg sc; Pfizer Inc) was administered, and animals were allowed to recover in thermostatically controlled cages. A post-operative period of two weeks was allowed before testing commenced.

O_2_ recordings were carried out on animals within standard mouse operant chambers (Med Associates, USA), where their head mounted 6-pin socket was connected to a low-noise, four-channel Quadstat (eDAQ Pty Ltd, New South Wales, Australia) via a flexible screened 6-core cable mounted through a swivel (both Plastics One Roanoke, VA) in the ceiling of the cage. This arrangement allowed free movement of the animal throughout the cage. The eCorder data acquisition system with Chart software (eDAQ Pty Ltd, Australia) were used for monitoring and recording O_2_ signals. As part of a larger study, a cohort of 22 animals were cabled for approximately 30 minutes pre-injection to establish a stable baseline. Animals were then treated with either vehicle or the relevant dose of xanomeline (1, 3, or 10mg/kg i.p.) and oxygen signals recorded for 90 minutes before they were returned to their home cage. Animals were dosed once a week in a randomised Latin square design so that all animals received all treatments by the conclusion of the experiment.

Amperometric recordings from each working electrode channel were recorded at 40 Hz, and linear interpolation was used to replace occasional missing data points and a bi-quad Butterworth filter (high-pass 0.1 Hz) was used to suppress fast noise-related artefacts. Data was normalized by subtraction of the 60 s average pre-dose value from each data point in the series, thereby compensating for absolute differences in baselines between channels. Finally, a boxcar-averaging algorithm was applied to down-sample the data, keeping a single average from multiple 120 s non-overlapping windows. For each regional signal, area under the curve (AUC) was calculated according to the formula 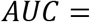 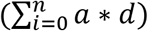, where n = number of samples in the curve, a = current (nA) at a given sample and d = sample interval (120 s). Statistics were conducted using Statistica v9 (StatSoft Inc, USA) software. AUC values were analyzed by a Repeated Measures ANOVA with region and dose as within-subject factors, followed by Fisher’s LSD post-hoc test for multiple comparisons. After exclusions for bad signal quality, statistical outliers and off-target histology placements, 6 animals were included in the dorsal striatum (DS) analysis and 6 animals included in the dorsal hippocampus (DHPC) analysis.

### EEG

Chronic measurement of EEG and electromyogram (EMG) was conducted using cranial implants placed under anesthesia previously described by (Seidel et al., 1995). The implant consisted of a miniature connector (Omnetics, USA) connected to five stainless steel screws positioned from bregma, with two frontal (+3.5mm AP, ± 2mm ML) and two occipital screws (−6.5mm AP, ± 5.2mm ML) for EEG recording and one overlying the cerebellum to be used as a ground. The implant housed two Teflon-coated stainless steel wires placed under each nuchal trapezoid muscle for EMG recordings. A miniature transmitter (Minimitter PDT4000G, Philips Respironics, Bend, OR) to monitor body temperature and locomotor activity was placed in the abdomen during the same surgery. Analgesics were used to minimize pain, buprenorphine (0.05mg/kg, SC) was administered pre-operatively, at the end of the surgery day and on the morning of the first post-operative day, and Metacam (meloxicam, 0.15 mg/kg, PO) was administered for 6 days after surgery. An antibiotic (Ceporex (cefalexin) 20 mg/kg PO) was administered 24 hours before and again immediately before surgery, and for 7 days after surgery. Rats were allowed to recover for at least 28 d prior to experimentation.

Animals individually housed within a specially modified polycarbonate cage with a flexible tether connecting the cranial implant to a commutator (Hypnion, USA). Each cage had individually controlled temperature and humidity with a strictly controlled 24-hour light-dark cycle (12h:12h, 5am lights on). Food and water were available ad libitum and consumption of each was measured via the break of infrared beams positioned in front of the food hopper and water bottle (Lixit). Drug treatments occurred 5 hours after lights on (CT-5), and rats were left undisturbed for 48 hours before and after each treatment with at least 6 days between treatments. Drug studies were conducted using a counterbalanced within-subject Latin-square design.

EEG signals, recorded as the differential between an ipsilateral pair (right hemisphere) of skull screws, were amplified 10,000x, bandpass filtered at 1-300Hz and digitized at 400Hz [Grass Corp., Quincy, MA]. EMG signals were amplified 20,000x, bandpass filtered at 10-100Hz and integrated based on the root mean square (RMS). EEG and EMG data were used in combination for on-line classification of arousal states into NREM sleep, REM sleep, or wake in 10s epochs. Wakefulness and sleep states were determined using SCORE-2006™ (Van Gelder et al., 1991), with 10 s epoch of EEG/EMG data classified as wake, NREM or REM sleep. Baseline wake, NREM sleep and REM sleep were calculated for 24 hours prior to dosing with post-treatment values calculated for 19 hours post treatment. Wake epochs were analyzed across multiple band frequencies including delta (0.1-4Hz), theta (4-9Hz), beta (12-30Hz), gamma (30-80Hz) and high frequency oscillation (130-160Hz) band power during the main treatment induced wake period. Quality control of the arousal state scoring was facilitated by visual assessment of the raw EEG and EMG signals by experts who were not involved in the data acquisition phase and were blinded with respect to treatment group. All drugs were made up fresh each testing day and administered at a final volume of 1ml/kg. Xanomeline was dosed at 1,3 and 10mg/kg by sub-cut injection on pH adjusted water. In study 2 xanomeline was dosed at 10mg/kg in pH adjusted water at CT-4.5, and ketamine sub-cutaneously at 10mg/kg in 5% glucose 30 minutes later.

A fast Fourier transform of the EEG signal produced a measure of the spectral power for each discreetly scored 10-second epoch and subsequently binned into the frequency bands: delta (0.1-4Hz), theta (5-9Hz), beta (12-30Hz), gamma (30-80Hz) and high frequency oscillations (130-160Hz). The effects of the compounds were evaluated over the first 7-hour light phase following treatment when the wake promoting effects were maximal. Statistical analyses of spectral parameters were performed using the SAS (version 9.2, SAS Institute, Inc., Cary, NC) software package. The mixed model procedure was used to perform an analysis of covariance with treatment. In addition, baseline characteristics (chambers, body weight, age, pre-treatment distributions) were checked for homogeneity across treatments and factors of Body Weight, Age, Pre*Dose interaction, and 2- and 3-way Run date interactions were also considered in the model for influence on final estimates. Data are displayed as means ± SEM, with all figures created using GraphPad Prism 6 (GraphPad Software, US). All tests of significance were performed with α=0.05.

---

**Supplementary Table 1:** Mean plasma concentration of xanomeline or PCP at the end of the fMRI experiments for the four experimental groups.

|  | Concentration Xan<br>(nM) | Concentration PCP<br>(nM) |
| --- | --- | --- |
| Xan – Veh | 686±34 | - |
| Xan – PCP | 1026±22 | 42±1.5 |
| Veh – PCP | - | 38±2 |

## Supplementary Figures

**Figure S1:**
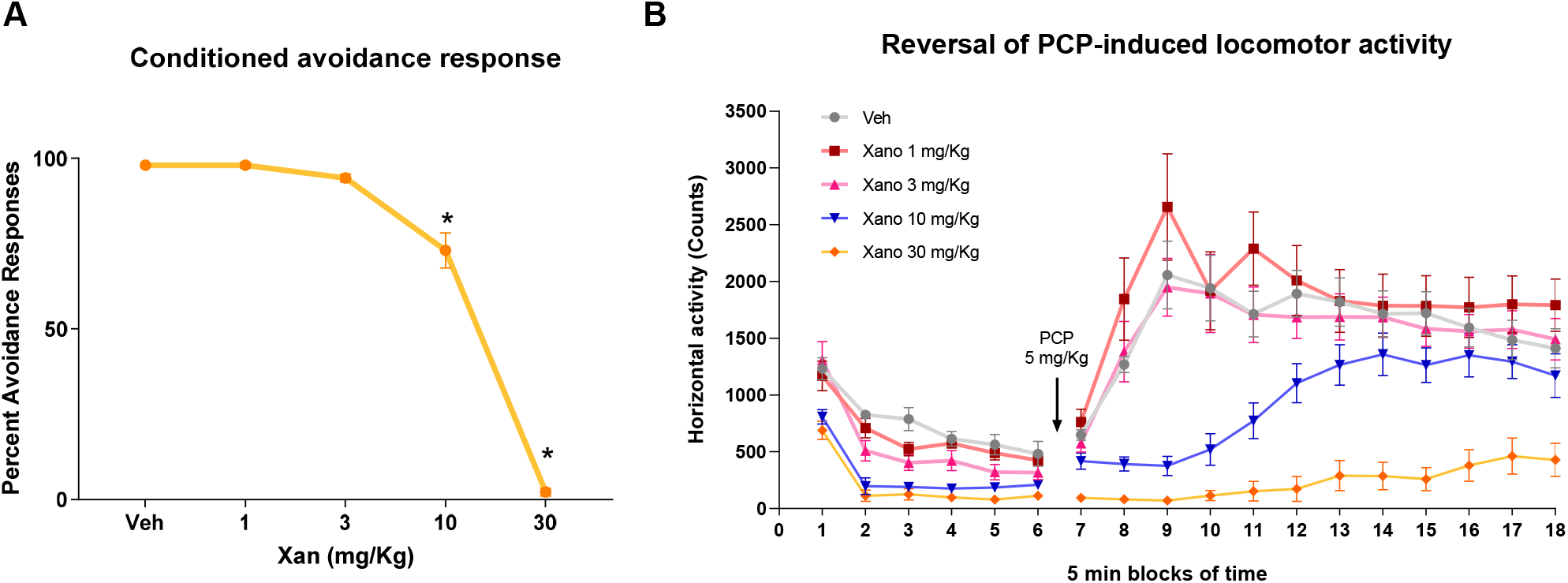
Behavioral determination of xanomeline dose. Dose-response curve of xanomeline tartrate. (A) Conditioned avoidance response (One-factor repeated-measures ANOVA with contrasts against the Saline control,*p < 0.0125, Bonferroni-corrected) and (B) reversal of PCP-induced locomotor activity assays. Total horizontal activity counts for each phase were analyzed using one-way ANOVA and Dunnett’s test, using the vehicle-treated group (prior to PCP-treatment) as the control for the habituation phase (spontaneous activity) and the vehicle-treated group (after PCP-treatment) as the control for the PCP-induced activity phase. In both (A and B), partial antipsychotic-like efficacy is observed at 10 mg/kg and complete efficacy at 30 mg/kg.

**Figure S2:**
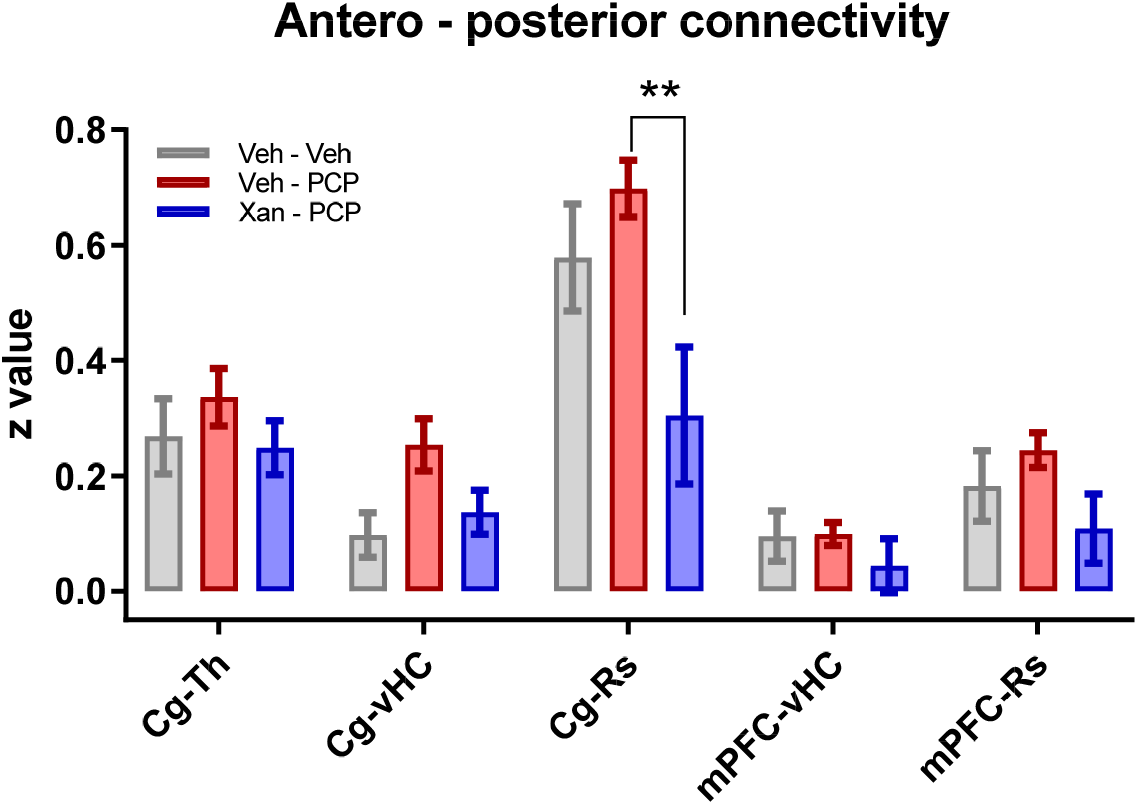
Xanomeline attenuates PCP-induced long-range functional connectivity changes. Quantification of antero-posterior rsfMRI connectivity (mean ± SEM, xan-PCP vs. veh-PCP. **p<0.01, student t test, FDR corrected). Abbreviation list: mPFC, medial prefrontal cortex; Cg, cingulate cortex; Rs, retrosplenial cortex; Th, thalamus; vHC, ventral hippocampus.

**Figure S3:**
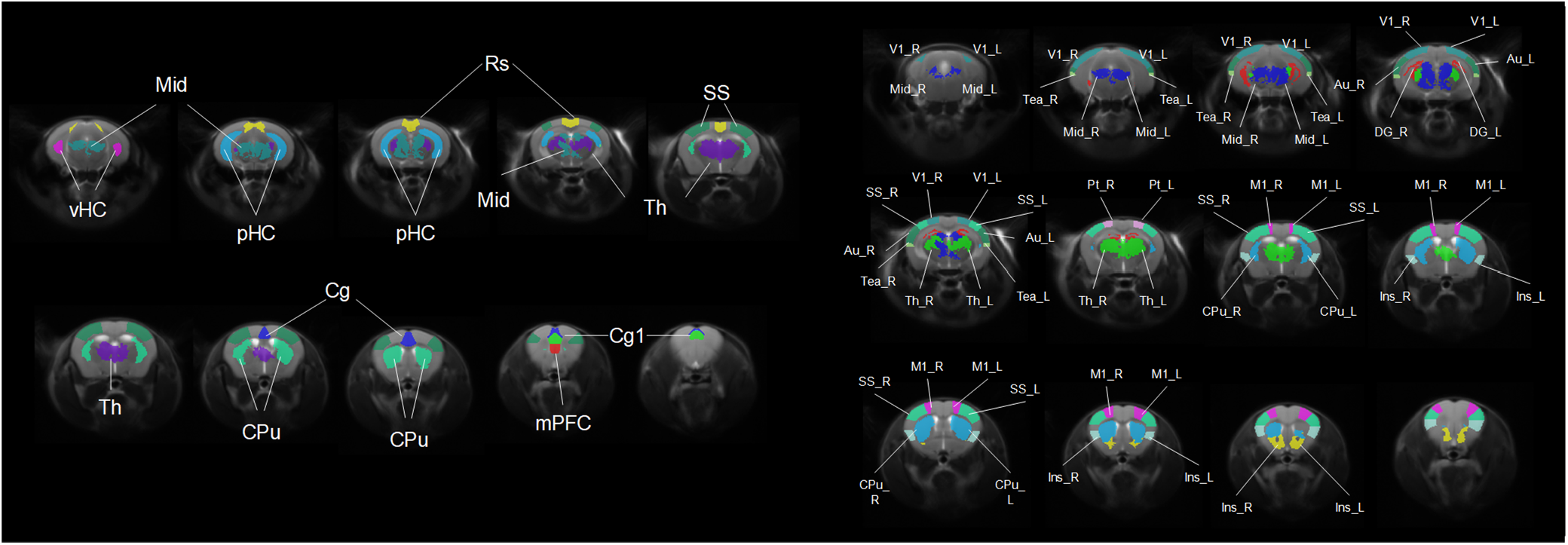
Anatomical location of (Left) bilateral and (right) mono-hemispheric volumes of interest employed for treatment quantification. Abbreviation list: mPFC, medial prefrontal cortex; Cg1, cingulate cortex; Cg, cingulate cortex; Rs, retrosplenial cortex; Th, thalamus; vHC, ventral hippocampus; pHC, posterior dentate gyrus; SS, primary, somatosensory cortex; CPu, striatum; Mid, midbrain; V1, primary visual cortex; Tea, temporal association cortex; DG, dentate gyrus; Au, auditory cortex; Pt, parietal cortex; M1, primary motor cortex; Ins, insula; L, left; R, right.

**Figure S4:**
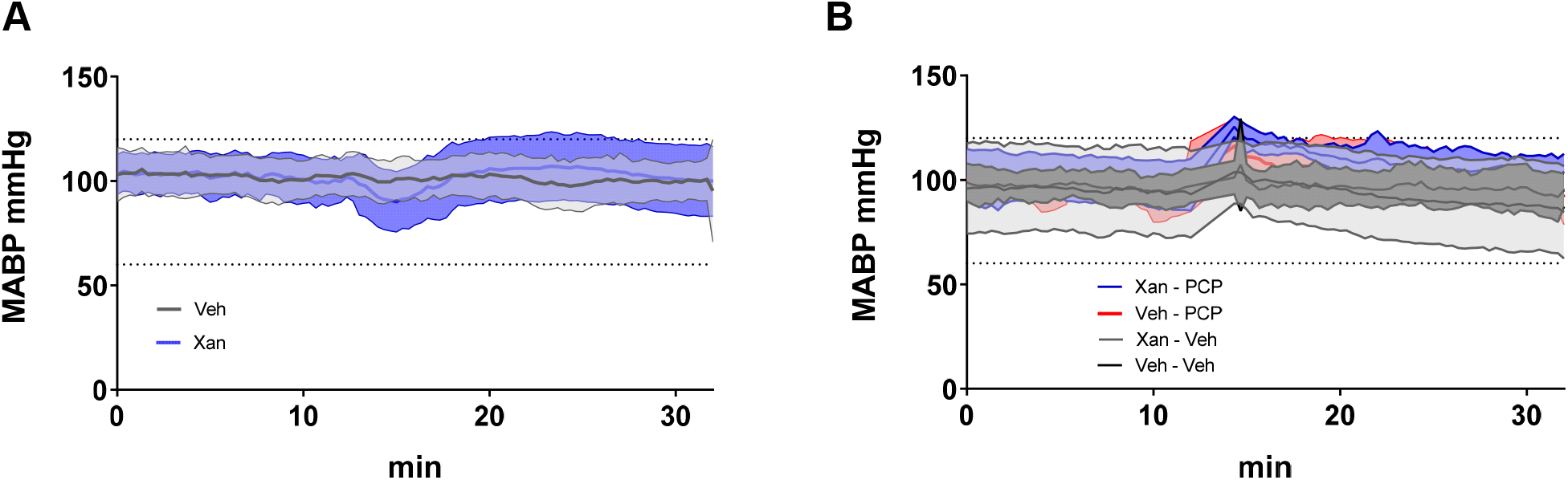
Mean arterial blood pressure (MABP) measured across treatment groups. (A) MABP changes produced by xanomeline (30 mg/Kg, n=18) or vehicle administration (n=19). (B) MABP changes produced by PCP (1mg/kg) administration in animals pretreated with xanomeline or vehicle. Xan – PCP, n=10; xan – veh, n=8; veh – PCP, n=10; veh – veh, n=9.

## Notes

### Competing Interest Statement

The authors have declared no competing interest.

